# Discovery of a new *Neisseria gonorrhoeae* Type IV pilus assembly factor, TfpC

**DOI:** 10.1101/2020.09.26.314724

**Authors:** Linda I. Hu, Shaohui Yin, Egon A. Ozer, Lee Sewell, Saima Rehman, James A Garnett, H Steven Seifert

**Affiliations:** Department of Microbiology-Immunology, Northwestern University Feinberg School of Medicine, Chicago, IL, USA; Division of infectious Diseases, Northwestern University Feinberg School of Medicine, Chicago, IL, USA; Centre for Host-Microbiome Interactions, Dental Institute, King’s College London, London, UK

## Abstract

*Neisseria gonorrhoeae* rely on Type IV pili (T4p) to promote colonization of their human host and to cause the sexually transmitted infection, gonorrhea. This organelle cycles through a process of extension and retraction back into the bacterial cell. Through a genetic screen, we identified the NGO0783 locus of *N. gonorrhoeae* strain FA1090 as containing a gene encoding a protein required to stabilize the Type IV pilus in its extended, non-retracted conformation. We have named the gene *tfpC* and the protein TfpC. Deletion of *tfpC* produces a nonpiliated colony morphology and immuno-transmission electron microscopy confirms that the pili are lost in the Δ*tfpC* mutant, although there is some pilin detected near the bacterial cell surface. A copy of the *tfpC* gene expressed from a *lac* promoter restores pilus expression and related phenotypes. A Δ*tfpC* mutant shows reduced levels of pilin protein, but complementation with a *tfpC* gene restored pilin to normal levels. Bioinformatic searches show there are orthologues in numerous bacteria species but not all Type IV pilin expressing bacteria contain orthologous genes. Co-evolution and NMR analysis indicates that TfpC contains an N-terminal transmembrane helix, a substantial extended/unstructured region and a highly charge C-terminal coiled-coil domain.

**Importance:** Most bacterial species express one or more extracellular organelles called pili/fimbriae that are required for many properties of each bacterial cell. The *Neisseria gonorrhoeae* Type IV pilus is a major virulence and colonization factor for the sexually transmitted infection, gonorrhea. We have discovered a new protein of *Neisseria gonorrhoeae* called TfpC that is required to maintain the Type IV pili on the bacterial cell surface. There are similar proteins found in the other members of the *Neisseria* genus and many other bacterial species important for human health.

## Introduction

*Neisseria gonorrhoeae* is the main causative agent of the sexually transmitted infection gonorrhea. There were 555,608 reported cases of gonorrhea reported in the US in 2017 and an estimated 86.9 million worldwide as well as an alarming rise in antibiotic resistance (1, 2). There are three major problems that complicate the treatment of gonorrhea. First, the rapid rise of antibiotic resistance has resulted in strains that are refractory to conventional treatments (3). Second, many patients are asymptomatic, remain untreated, and contribute to spread of the disease. Third, infection does not result in long-term immunity to reinfection. These attributes have made a vaccine or novel antimicrobials desirable, but to date there are no viable novel treatments. The uncertainty for future treatment options emphasizes the need for new knowledge about *N. gonorrhoeae* colonization, pathogenesis, and innovative modes of treatment.

Almost all Gram-negative bacteria and a subset of Gram-positive bacteria express T4p (4). There are three major subsets of T4p, and there is a clear evolutionarily relationship with Type II secretion system (T2S) and Archaeal flagella (5). T4p provide a wide range of phenotypes to the organisms that express them and are important organelles that promote bacterial colonization and pathogenesis. The *N. gonorrhoeae* T4p is the only known virulence factor absolutely required for colonization (6-8). The type I T4p of *Neisseria meningitidis* are closely related to the *N. gonorrhoeae* T4p and are also necessary for colonization and disease (9).

The T4p has multiple functions that are critical for *N. gonorrhoeae* pathogenesis. The pilus is an essential factor for colonization, enhancing the ability of the bacterium to adhere to and interact with host cells and tissues at infection sites (10). The pilus is also required to promote bacteria-bacteria interactions, the formation, and dissolution, of microcolonies and biofilms (11). The T4p is required for twitching motility, a specialized form of locomotion that requires T4p retraction (9), to enhance bacterial interactions with the epithelium (12). The T4p apparatus is a bidirectional secretion apparatus that engages in pilus secretion, importing DNA for genetic transformation and the spread of antibiotic resistance, importing the pilus for twitching motility, and importing other molecules like antibiotics (13). T4p expression greatly increases Gc resistance to the oxidative and non-oxidative killing mechanisms of PMNs (14). While we have a grasp of many molecular mechanisms underlying *Neisseria* T4p assembly and function, many questions remain about how this dynamic fiber functions in pathogenesis.

Several proteins are involved in the assembly and function of the T4p. The main pilin subunit PilE starts as a prepilin with a 7 amino acid (AA) leader sequence. After secretion through the inner membrane (IM), the leader sequence is cleaved by the PilD signal peptidase to produce the mature protein (15). PilD is also required to process the minor pilins that share the N-terminal AA sequence similarity with pilin (16, 17). The PilQ protein is of the secretin class and forms a pore through the outer membrane (18, 19). The PilC protein (20) is localized to the outer membrane and has been implicated in contributing to adherence and modulating pilus retraction (21, 22). PilC is also reported as being localized to the pilus tip (23). PilP and PilW have been shown to interact with PilQ (24). The minor pilin proteins, PilH-L, are proposed to prime pilus assembly within the periplasm (25). The minor pilins PilV and ComP are dispensable for pilus assembly but have specific roles in adherence and transformation (16, 17). The PilF (aka PilB), PilT, and PilU proteins are cytoplasmic NTPases involved in modulating pilus extension and retraction (15, 26, 27).

We previously demonstrated that the activity of the Mpg zinc-metalloprotease is required to maintain T4p exposed on the bacterial cell surface (14). We also showed that Mpg activity on the T4p mediates protection from both oxidative and non-oxidative killing mechanisms of PMNs (14, 28). The increased sensitivity of nonpiliated cells to oxidative and nonoxidative killing is phenocopied by nonpiliated *N. gonorrhoeae* sensitivity to the iron-dependent, antimicrobial compound streptonigrin (SNG). A transposon sequencing (InSeq) screen for mutants that alter SNG sensitivity revealed a new T4p assembly factor we named TfpC.

## Results

We conducted a saturating, InSeq screen of Gc strain FA1090 to identify genes that when inactivated provided decreased or increased survival to SNG lethality. We will report the rationale and full results of the InSeq SNG screen elsewhere. From this screen, we identified the NGO0783 locus of *N. gonorrhoeae* strain FA1090 as providing an average 38.7-fold decrease in representation under 0.4 μM SNG selection when interrupted with any of nine distinct transposon insertion sites within the open reading frame.

We constructed a loss-of-function mutant of the NGO0783 locus (Figure 1A) in strain FA1090 by deleting most of the open reading frame and inserting a nonpolar (Table 1), kanamycin resistance gene (KmR). The Δ*tfpC* mutant showed a characteristic P-colony morphology, with flatter colonies with more spreading that results in larger diameter colonies than the P+ parent without the P+ dark edge (Figure 2A) (6). These changes in colony morphology are consistent with a reduction in pilus expression (29) or an effect on pilus bundling (30). We introduced the Δ*tfpC* mutation into three other *N. gonorrhoeae* isolates, and in each strain, this mutation resulted in P-colony morphology (Figure 2B). The mutant also showed reduced transformation competence (5.6 x 10^−5^ transformants/CFU vs 1.3 × 10^−3^ transformants/CFU for the piliated parent), consistent with the nonpiliated colony morphology, but the NGO0783 mutant is more competent for DNA transformation than a Δ*pilE* mutant that does not transform under these conditions (< 8 × 10^−7^ transformants/cfu). This phenotype is similar to other *N. gonorrhoeae* mutants that can still assemble pili but cannot maintain them in an extended conformation (14, 25, 31). However, in contrast to those mutants, introducing a *pilT* loss-of-function mutation to the NGO0783 mutant did not restore the parental P+ colony morphology (Figure 2B).

**Table 1:**
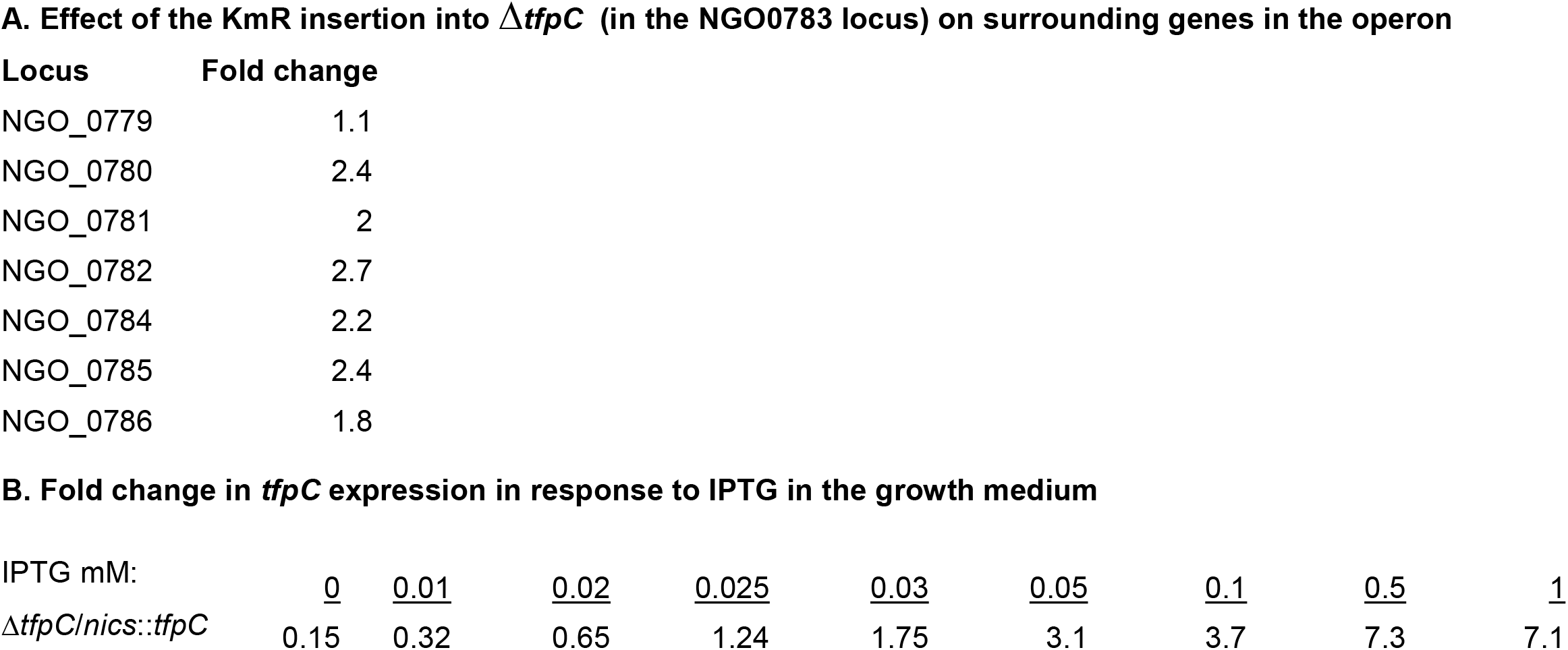
Quantitative RT-PCR

**A. Effect of the KmR insertion into $\Delta tfpC$ (in the NGO0783 locus) on surrounding genes in the operon**
| Locus | Fold change |
| --- | --- |
| NGO_0779 | 1.1 |
| NGO_0780 | 2.4 |
| NGO_0781 | 2 |
| NGO_0782 | 2.7 |
| NGO_0784 | 2.2 |
| NGO_0785 | 2.4 |
| NGO_0786 | 1.8 |

**B. Fold change in *tfpC* expression in response to IPTG in the growth medium**
|  |  |  |  |  |  |  |  |  |  |
| --- | --- | --- | --- | --- | --- | --- | --- | --- | --- |
| IPTG mM: | <u>0</u> | <u>0.01</u> | <u>0.02</u> | <u>0.025</u> | <u>0.03</u> | <u>0.05</u> | <u>0.1</u> | <u>0.5</u> | <u>1</u> |
| $\Delta tfpC/nics::tfpC$ | 0.15 | 0.32 | 0.65 | 1.24 | 1.75 | 3.1 | 3.7 | 7.3 | 7.1 |

**Figure 1.**
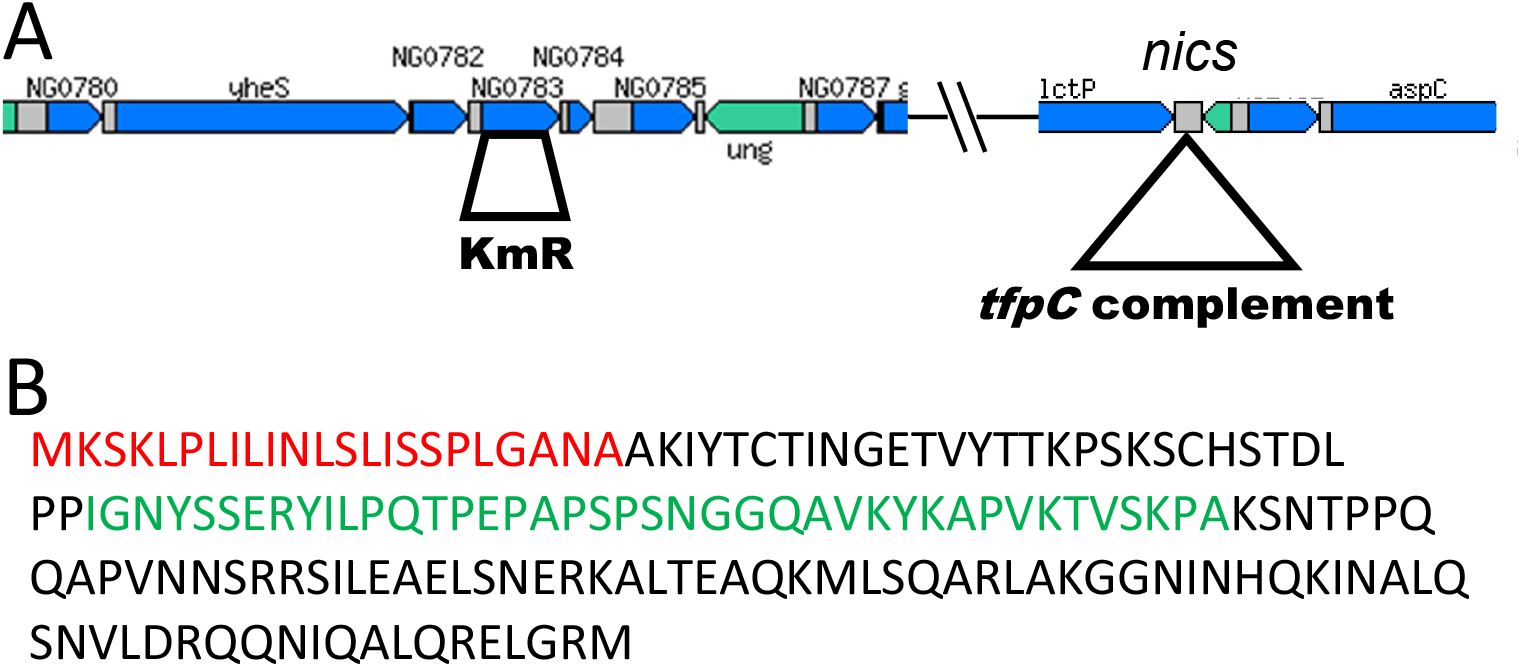
Cartoon of the NGO0783 locus, mutants and complements. A. Cartoon of the chromosomal regions of strain FA1090 with the NGO0783 locus and the surrounding loci and the location of the *nics* chromosomal complementation site (59) expressing the *tfpC::flag* complement (from: http://stdgen.northwestern.edu). B. Predicted amino acid sequence of the *N. gonorrhoeae* TfpC protein. The cleavable signal sequence is shown in red.

**Figure 2:**
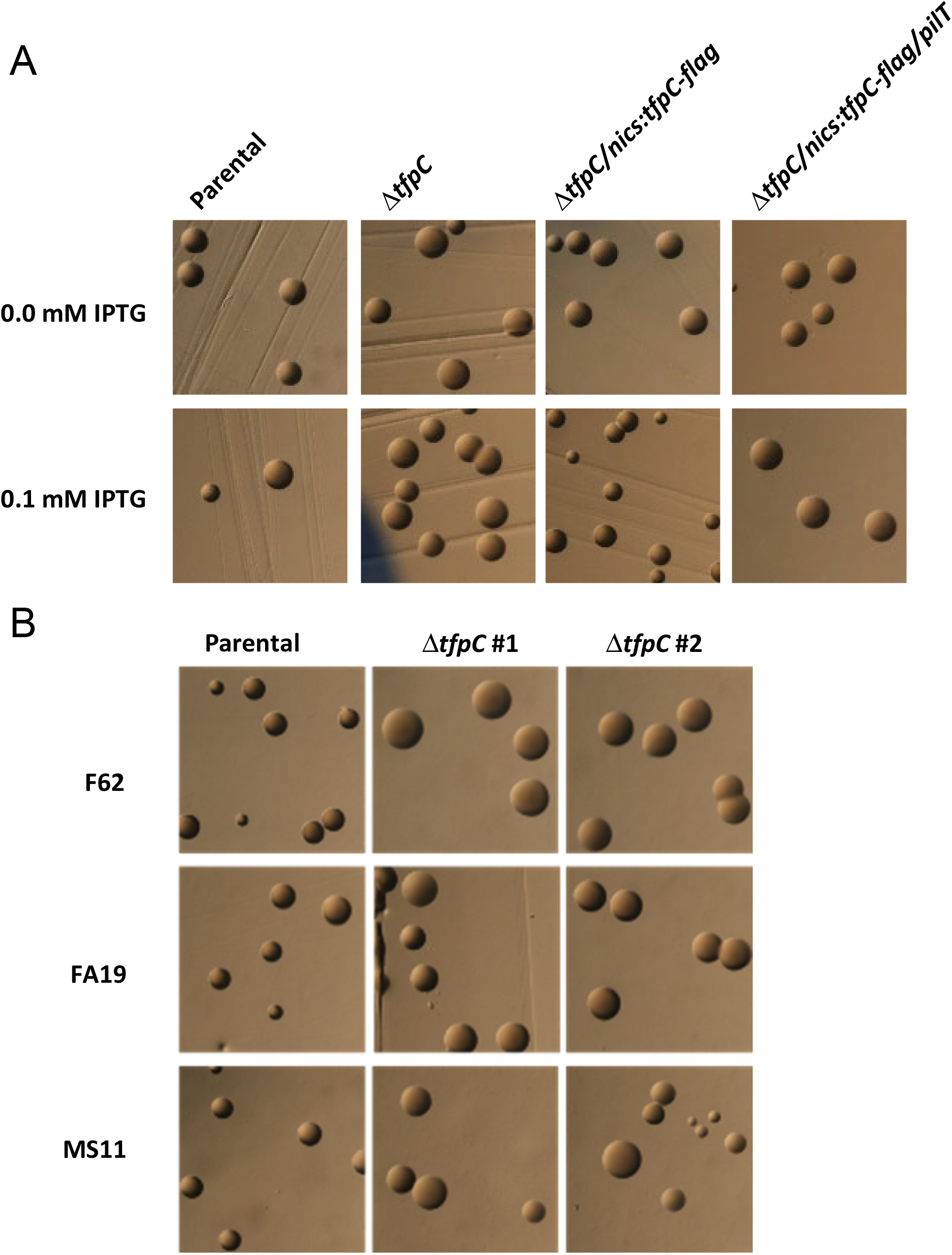
Analysis of Pilus-dependent Colony Morphology. A. Stereo-micrograph pictures of 22 hr *N. gonorrhoeae* colonies grown with or without 0.1 mM IPTG in the medium. The parental strain is FA1090 1-81-S2 *recA6*. B. Pilus dependent colony morphology changes in two different Δ*tfpC* transformants of *N. gonorrhoeae* strains F62 FA19, and MS11 grown for 22 hrs. Piliated colonies (e.g., Parental strain) are smaller, have a dark ring at the edge of the colony and are domed. Nonpiliated colonies (e.g., Δ*tfpC* strain) are larger, have no dark ring or a less pronounced ring, and are flatter.

We introduced a series of IPTG-regulated, complementation constructs at an ectopic locus in the FA1090 chromosome to express native TfpC, as well as an epitope tagged version (FLAG-tagged) (Figure 1A). The IPTG-regulated *tfpC-flag* complement construct restored a piliated colony morphology (Figure 2A) and SNG resistance (not shown). We used the Flag-tagged complement for all further analysis. Based on these preliminary results, we predict that this gene is involved in T4p elaboration, and therefore named the gene within the NGO0783 locus as *tfpC* for <u>T</u>ype <u>f</u>our <u>p</u>ilus assembly protein <u>C</u>, and the protein as TfpC. We cannot rule out other roles for the TfpC protein in cellular processes distinct from piliation, but did not observe any obvious cellular phenotypes that would suggest an alternative function.

Wild type levels of *tfpC* mRNA were produced from the complemented strain when 0.025 mM IPTG was added to the growth medium (Table 1B). Western blot analysis of TfpC protein levels confirmed the Q-RT-RCR results (Figure 3A). Analysis of total pilin levels with pilin tagged with a C-Myc epitope tag (32) showed that the Δ*tfpC* mutant had lower levels of pilin protein compared to the parental FA1090 strain that were restored when the complemented strains was grown with 0.025 mM IPTG in the growth medium (Figure 3B). Moreover, 0.1 mM IPTG produced 3.7-fold higher levels of *tfpC* mRNA (Table 1). Even with overexpression of *tfpC*, there was no noticeable growth (data not shown) or colony morphology phenotype (Figure 2A) compared to the parental strain. When we grew the complemented strain with 0.1 mM IPTG in the medium, there was a small increase in the pilin protein band relative to the parental strain. Surprisingly, a Δ*tfpC/*Δ*pilT* double mutant had more total pilin protein by Western blot than the Δ*tfpC* mutant alone. These results show that TfpC acts to stabilize pilin, the loss of the pilus in the Δ*tfpC* mutant is dependent on PilT, and the absence of pilus retraction stabilizes pilin in both wild-type TfpC expressing strains and the Δ*tfpC* mutant background.

**Figure 3:**
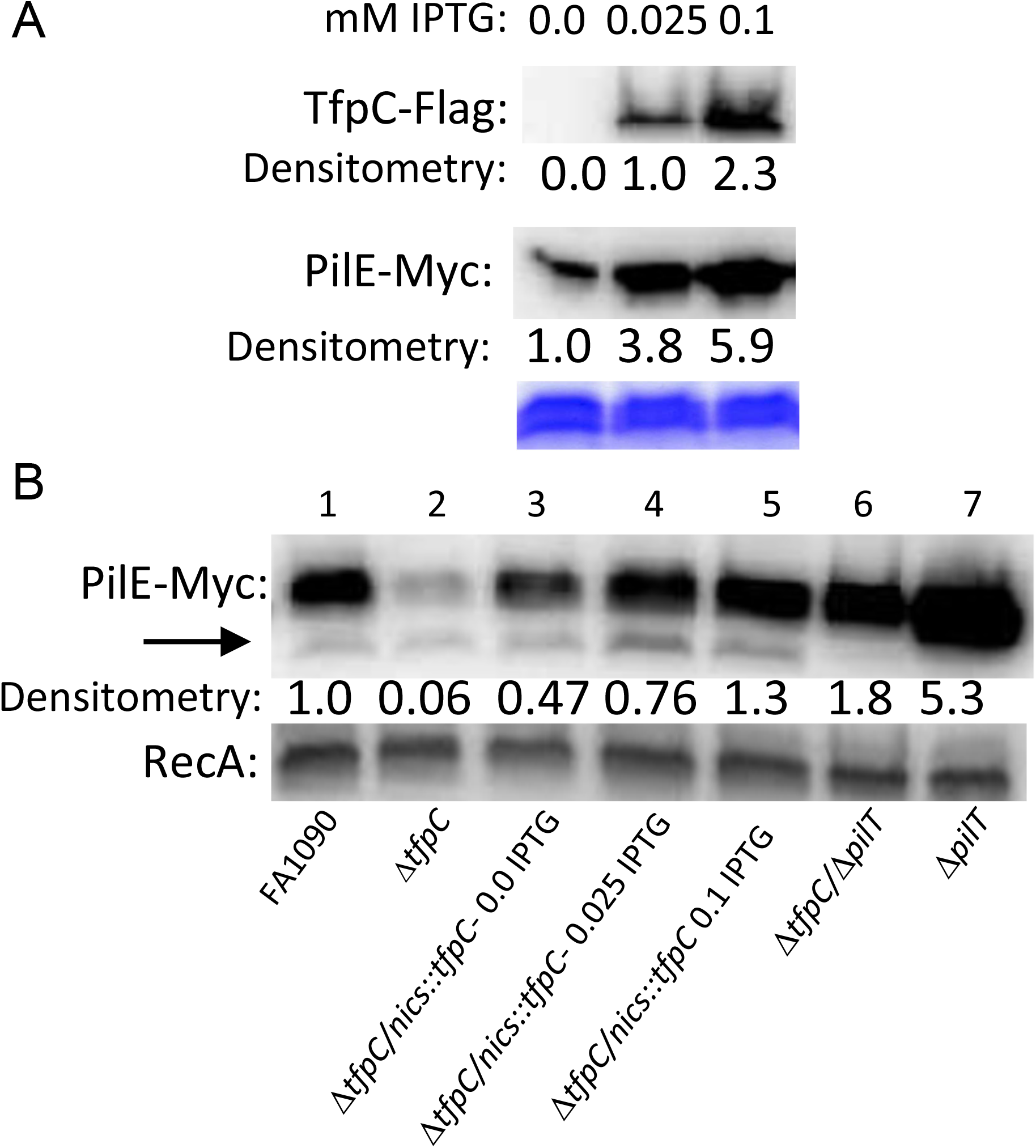
Western Blot Analysis of total TfpC and Pilin expression. A. Strain FA1090 1-81-S2 *pilE-myc* Δ*tfpC/nics*::*tfpC-flag* was grown with different levels of IPTG probed and whole cell lysates were probed with with anti-FLAG Mab or anti-Myc-MAb. The section of the Coomassie stained gel shows equal loading of the proteins in the replicate gels. Estimates of relative protein amounts as determined by Densitometry are shown below each blot. Representative Western blot of three independent repeats. B. Strains Δ*tfpC*, Δ*tfpC/nics*::*tfpC-flag*, Δ*tfpC*/Δ*pilT*, and Δ*pilT* in FA1090 1-81-S2 myc-tagged *pilE* background (Q155) were grown with different levels of IPTG and whole cell lysates were probed with anti-Myc Mab. After development, the blot was washed and reprobed with anti-RecA antisera (*E*. coli). Western blot of two independent repeats. We presume that the smaller band indicated by the arrow is the truncated pilin form S-pilin (60).

We determined the effect of the Δ*tfpC* mutation on piliation in FA1090 using the C-Myc epitope tagged *pilE* (32) to allow visualization of the pilus using gold-labeled, secondary antibody by immuno-transmission electron microscopy (IM-TEM) (Figure 4 and Figure 5). These type of electron micrographs have limitations since they cannot quantitate pili because the bacterial cells are absorbed from a bacterial colony onto the grid. However, the IM-TEMs showed that the Δ*tfpC* mutant lost piliation (Figure 4) and that the Flag-tagged *tfpC* restored the pilus in the complemented strain (Figure 5). These results were consistent with the colony morphology phenotypes. Interestingly, many of the Δ*tfpC* mutant cells still had antibody binding near the bacterial cell surface (Figure 4), which was not observed with a non-piliated, *pilE* mutant strain. This result suggested that there were short pili on the cell surface or another form of pilin that reacts with the antibody near the cell surface. The IM-TEM analysis of the Δ*tfpC/*Δ*pilT* double mutant showed that loss of PilT restored pilus expression to the Δ*tfpC* mutant (Figure 5), a result consistent with the Western blot analyses (Figure 3). However, the pili in the Δ*tfpC/*Δ*pilT* double mutant did not show any essential differences from those expressed on the parental strain.

**Figure 4:**
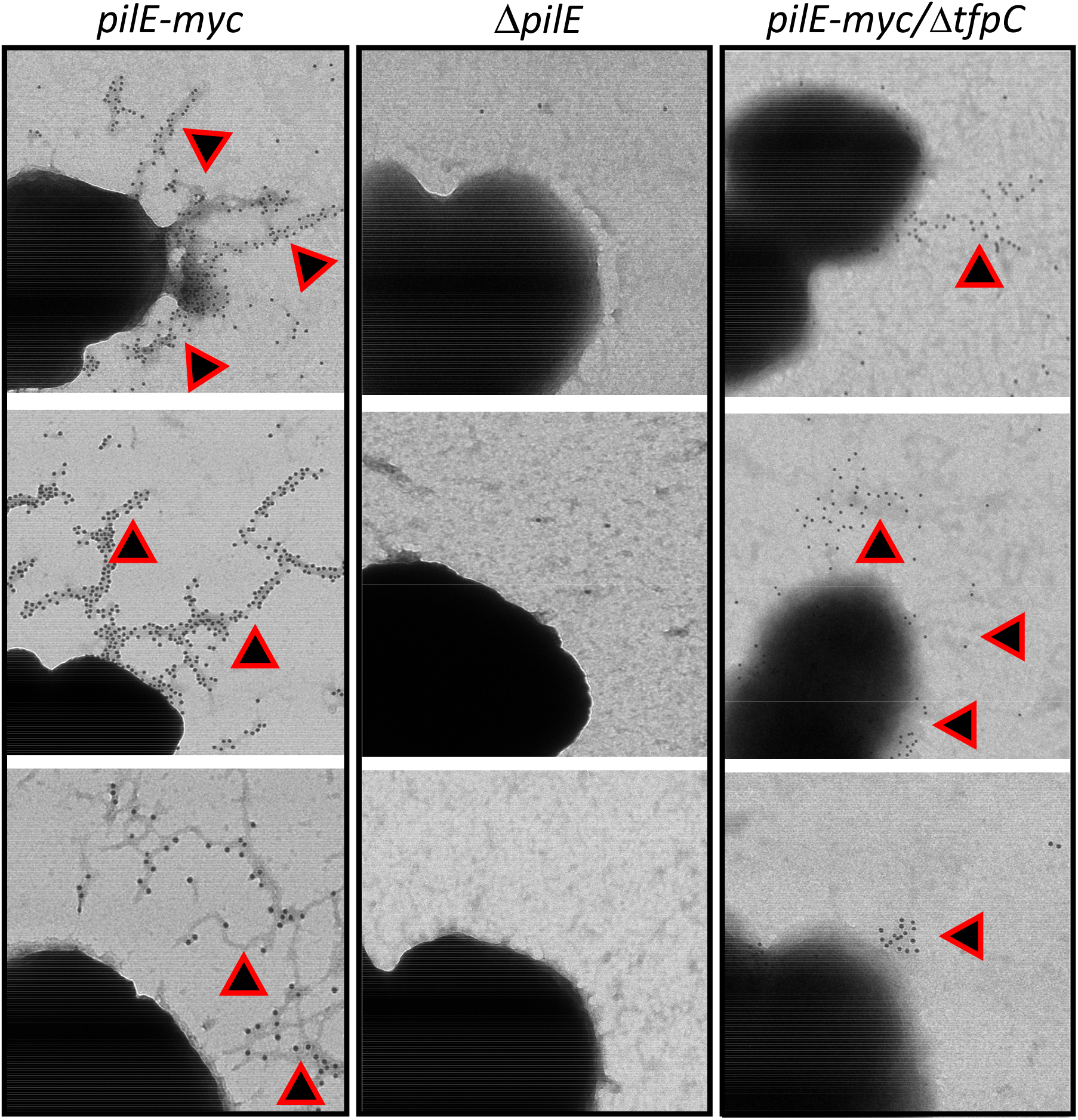
Immuno-TEM Micrographs of Pilus expression on Δ*tfpC* Mutant. Micrographs of cells lifted onto grids from 22 hr colonies of FA1090, *pilE-myc* strain; FA1090, Δ*pilE* nonpiliated mutant; and FA1090, *pilE-my,c* Δ*tfpC* mutant that were reacted with anti-Mac-Mab and then secondary gold-labeled, anti-mouse IgG. The small round gold particles show were immune-reactive pilin is localized and are highlighted with triangles. These are representative images from two independent experiments.

**Figure 5:**
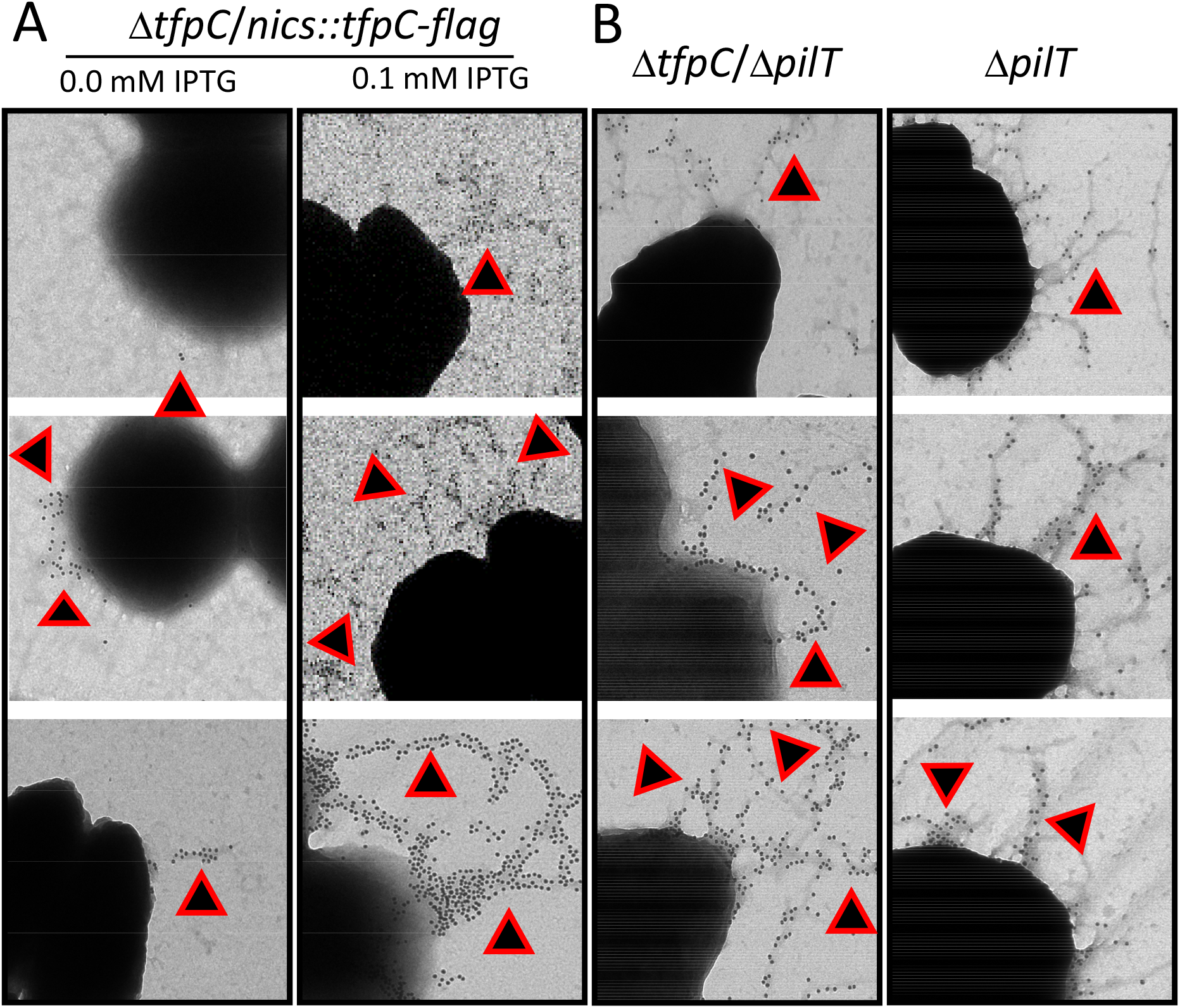
Immuno-TEM of Pilus expression. Representative micrographs of cells lifted onto grids from 22 hr colonies of: A. FA1090 *pilE-myc* strain, Δ*tfpC*/*nics::tfpC-flag* grown with or without IPGT to induce TfpC expression. FA1090 *pilE-myc*, Δ*<u>tfpC</u>*, Δ*pilT* and FA1090 *pilE-myc*, Δ*pilT*. that were reacted with anti-Mac-Mab and then secondary gold-labeled, anti-mouse IgG. The small round gold particles show were immune-reactive pilin is localized and are highlighted with triangles. These are representative images from two independent experiments.

**Figure 6:**
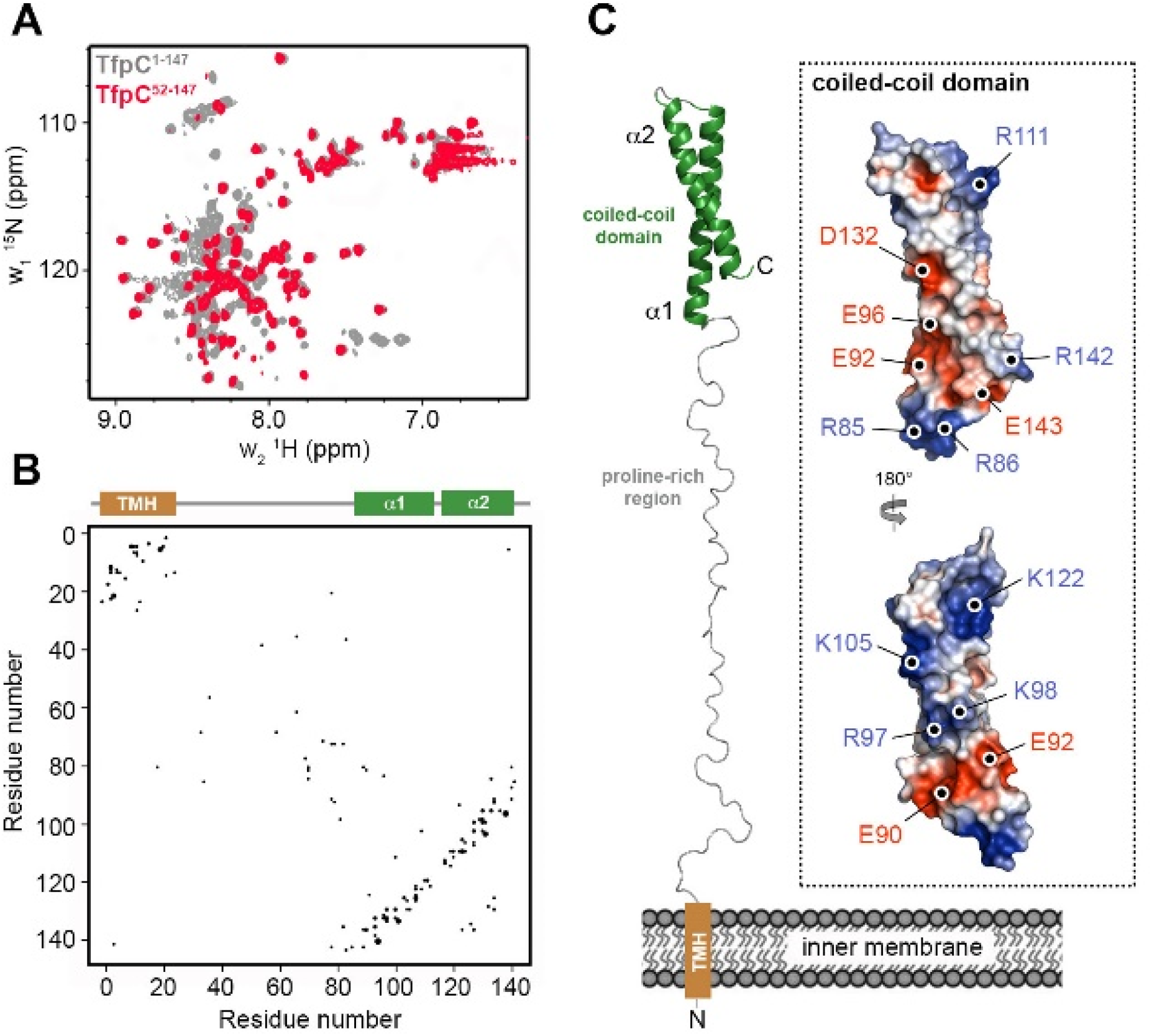
Structural Model of TfpC. (A) Overlay of ^1^H-^15^N HSQC NMR spectra for mature TfpC (residues 1-147; grey) and N-terminally truncated TfpC (residues 52-147; red). Proton resonances observed between ∼ 8.0 and 8.5 ppm indicate the presence of characteristic clusters of unstructured backbone amides. The peaks resonating at high chemical shift (>8.5 ppm) correspond to highly ordered backbone amides present in secondary structure elements. However, lack of dispersion (no peaks >9.0 ppm) suggests the presence of an extended helix or coiled-coil structure. Removal of N-terminal residues from TfpC results in a reduction of disordered resonances. (B) Co-evolution contact map for TfpC with secondary structure features highlighted. The transmembrane helix (TMH) is brown and helices are green. (C) Co-evolved coupling restrained model of TfpC. Inset shows the C-terminal coiled-coil domain as electrostatic surface potential, with charged surface residues highlighted. The surface of the C-terminal domain is composed of both large positive and negative patches, which may mediate recognition of partner protein(s).

Bioinformatic analysis indicated that TfpC has a cleavable periplasmic localization signal at its N-terminus, followed by a short transmembrane helix, an extended proline-rich region, and a helical domain at the C-terminus (Figure 1B). This predicted structure was supported by NMR experiments where we compared ^1^H-^15^N HSQC spectra for mature recombinant TfpC (residues 1 to 147; minus the signal sequence) and an N-terminally truncated TfpC (residues 52 to 147) (Figure 5A). Proton resonances for the N-terminal region of TfpC were observed between ∼ 8.0 and 8.5 ppm, indicative of unstructured peptide, while highly ordered backbone amides peaks (>8.5 ppm) did not extend above 9.0 ppm, which suggested the presence of an extended helix or coiled-coil structure at the C-terminus.

Analysis of the co-evolution between different amino acid sites within a protein sequence can provide strong evidence for inter-residue interactions, such as those found in protein sub-domains (33). We therefore performed co-evolution analysis on the mature TfpC sequence using the EVcouplings Python framework (34). We identified 1065 similar sequences and used in the alignment stage, which provided an excellent alignment solution with a ratio of effective sequences to protein length of 5.15 (Figure 5B). There were 77 strong evolutionary couplings identified, which generally clustered between residues located either in the transmembrane helix region or the C-terminal helical domain, but very few couplings were observed in the extended central region. WE tehn used these couplings as distance restraints to generate a model of TfpC. The model suggests that the N-terminus of TfpC may insert into the bacterial inner membrane, while a C-terminal coiled-coil domain is projected into the periplasm via an extended proline-rich region (Figure 5C). The C-terminal domain contains a high proportion of charged residues and the surface of the model is composed of both large positive and negative patches. This indicates that this region may be involved in the recognition of partner protein(s), presumably in the periplasm, and that electrostatic interactions drives important interactions with the pilus machinery (Figure 5C).

## Discussion

Almost every Gram-negative bacterial species expresses at least one Tfp and as do many Gram-positive organisms. The apparatus that allows the expression and function of the T4p spans the bacterial envelope and is evolutionarily related to the T2S apparatus in many bacterial species and the archaeal flagella. The assembly and function of these organelles have been studied intensively in these disparate types of prokaryotes. Since the InSeq transposon screen was to identify gene products involved in resistance or sensitivity to streptonigrin, we were intrigued when we found that the NGO0783 locus contained a gene product important for piliation.

Bioinformatic analysis of the open reading frame in the NGO0783 locus provided several predictions about the protein structure and function. The predicted protein has a predicted molecular weight of 18.455 kd and a basic pI of 10.8. The TfpC protein has a standard, Sec-dependent, cleavable signal sequence (Probability=0.99 by SignalP 5.0) and the mature protein has a hydrophobic N-terminus with many proline residues (Figure 1B). The best-fit structural prediction model from Phyr2 is a HR1 repeat protein with regions connected by a central hinge (**Error! Reference source not found**.Figure 5). There is enrichment of the TfpC protein in cell envelopes and membrane vesicles when the MlaA phospholipid removal protein is inactivated (35) showing TfpC is localized to the bacterial envelope. The TfpC protein sequence is 99-100% conserved in all sequenced *N. gonorrhoeae* isolates, suggesting it is not surface exposed. Based on these analyses we predict that this ORF localizes to the bacterial periplasm.

BLASTP revealed that the ORF has a DUF4124/pfam13511 domain of unknown function and is the only member of the cl16293 superfamily of proteins. There are orthologues of TfpC present in genomic sequences of *Neisseria meningitidis, Neisseria lactamica, Neisseria polysaccharea* and *Neisseria cinerea* with 100% amino acid identity. There were also other *Neisseria sp*. orthologues with lower but significant similarity, including several with an N-terminal extension and an additional middle domain not found in in the *N. gonorrhoeae* orthologue. A search of the Pfam database shows 948 bacterial species in many genera with proteins with the DF13511 (DUF4124) domain however how many of these proteins are true orthologues and involved in T4p or T2S is not known from this type of analysis. In our searches, we found the Dsui_1049 locus of the bacterium *Dechlorosoma suillum PS* (an environmental Gram-negative also called *<u>Azospira oryzae</u>* (36)) that shows a Waterman_Eggert score of 169 and E<5.6e-10 with TfpC. Many of the genes in *D. suillum PS* that show a fitness correlation with a Dsui_1049 mutant are T4p-associated genes, supporting a broad role for TfpC orthologues in piliation (Fitness Browser - http://fit.genomics.lbl.gov). Interestingly, there are other well-studied T4p-expressing species with no close orthologue, such as *Pseudomonas aeruginosa* and *Vibrio cholerae*. These species do have DUF4124 domain proteins but there is too limited sequence similarity to assign these as orthologues. It will be interesting to determine why only some species that express T4p have a TfpC orthologue and whether the more distant orthologues are all involved in T4p expression or could alternatively be involved in T2S or other related processes.

Introducing a Δ*pilT* loss-of-function mutation into the Δ*tpfC* strain produced two contrasting phenotypes. The inactivation of pilus retraction through loss of PilT did not restore the piliated colony morphology, but the TEMs clearly showed that pili where restored when PilT was inactivated and there was no observable difference between the parental pili and the pili observed with the Δ*tpfC/*Δ*pilT* mutant. We assume that the pili expressed on the Δ*tpfC/*Δ*pilT* are different in a way that alters the colony morphology hat is not reflected in the TEMs.

One of the more interesting phenotypes of the Δ*tfpC* mutant is the loss of the pilin protein in the mutant and the stabilization of pilin when we overexpressed TfpC with 0.1 mM IPTG (Figure 3). The observation that loss of pilus retraction in the Δ*tfpC/*Δ*pilT* double mutant also stabilizes pilin suggests that the role of TfpC in stabilizing pilin occurs after pilus retraction and not during pilus extension or within the extended fiber. However, the fact that a Δ*pilT* mutant strain with wild-type *tfpC* also shows a stronger pilin band suggests that PilT-dependent pilin degradation occurs all the time. This observation of a retraction-dependent destabilization of pilin has been previously reported for strain MS11 (37). We propose that pilin that is within the assembled pilus fiber is protected from proteolysis, but that upon retraction pilin becomes exposed to periplasmic proteases. In the future, determining whether proteolysis occurs during the process of retraction or after pilin returns to the cytoplasmic membrane will provide important insight into T4p dynamics.

Based on the phenotypes of the Δ*tfpC* mutant and Δ*tfpC/*Δ*pilT* double mutant, we propose that the TfpC protein is not necessary for T4p expression but rather is necessary to maintain the T4p in an extended state until retraction occurs. We speculate that the surface-associated pilin detected in the Δ*tfpC* mutant (Figure 4) could be pili caught in the process of retraction. This same PilT-dependent loss of pilus expression occurs when several other pilus-associated proteins are inactivated and we have proposed that there might be a peptidoglycan linked anti-retraction complex that mediates this phenotype since mutants lacking several peptidoglycan modifying enzymes (Mpg and DacB/C) also show a PilT-dependent modulation of pilus expression (14, 38). There are other mechanisms that could account for this phenotype, such as a role of TfpC and other proteins in modulating PilT activity, acting through the inner membrane complex. Determination of the precise subcellular localization of TfpC and its interaction partners will be required in future work to devise the mechanisms of TfpC in modulating pilus dynamics.

## Materials and Methods

### Strains and Growth

The studies performed here mainly used *N. gonorrhoeae* strain FA1090 PilE variant 1-81-S2 (39) and its isogenic derivatives. Strains MS11, F62 and FA19 were also tested (Table 2). The sequence of *pilE* was confirmed to be 1-81-S2 using PCR and sequencing with primers pilRBS and SP3A (Table 4). *N. gonorrhoeae* were grown in GC Medium Base (Difco) plus Kellogg supplements I and II [22.2 mM glucose, 0.68 glutamine, 0.45 mM cocarboxylase, 1.23 mM Fe(NO3)3] (GCB) at 37 °C in 5% CO_2_. Antibiotics and their concentrations used for selection in GCB were kanamycin (Kan) 50 μg/ml and erythromycin (Erm) 2 μg/ml. *E. coli* strains One Shot TOP10 Electrocomp *E. coli* (Invitrogen), DH5α, and BL21 (DE3) (New England Biolabs) used to propagate plasmids or protein were grown in Luria-Bertani (LB) solid containing 15 g/L agar or liquid media at 37 °C. The antibiotics and their concentrations used in LB were ampicillin (Amp) 100 μg/ml.

**Table 2:**
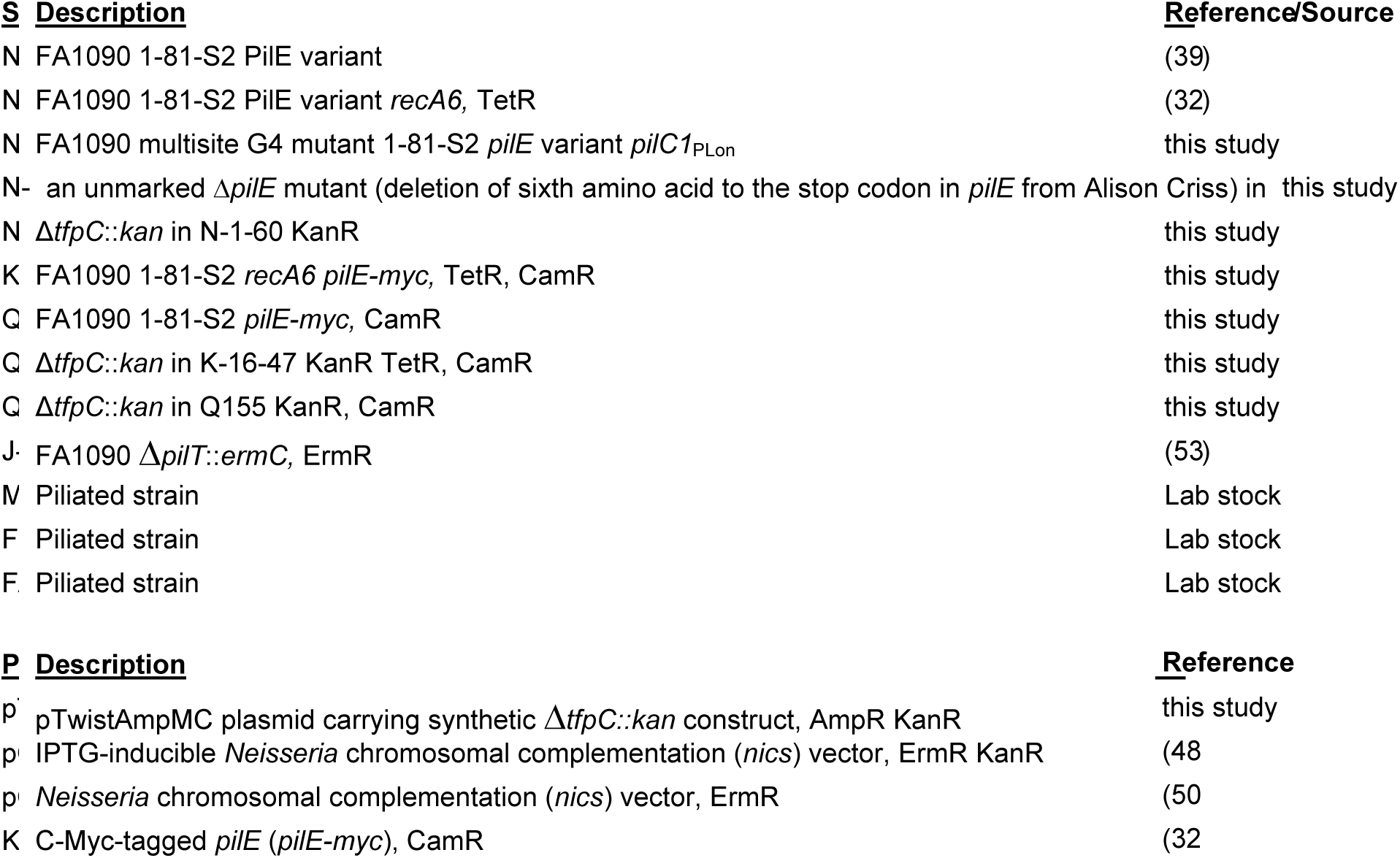
*N. gonorrhoeae* Strains and Plasmids

**Table 3.**
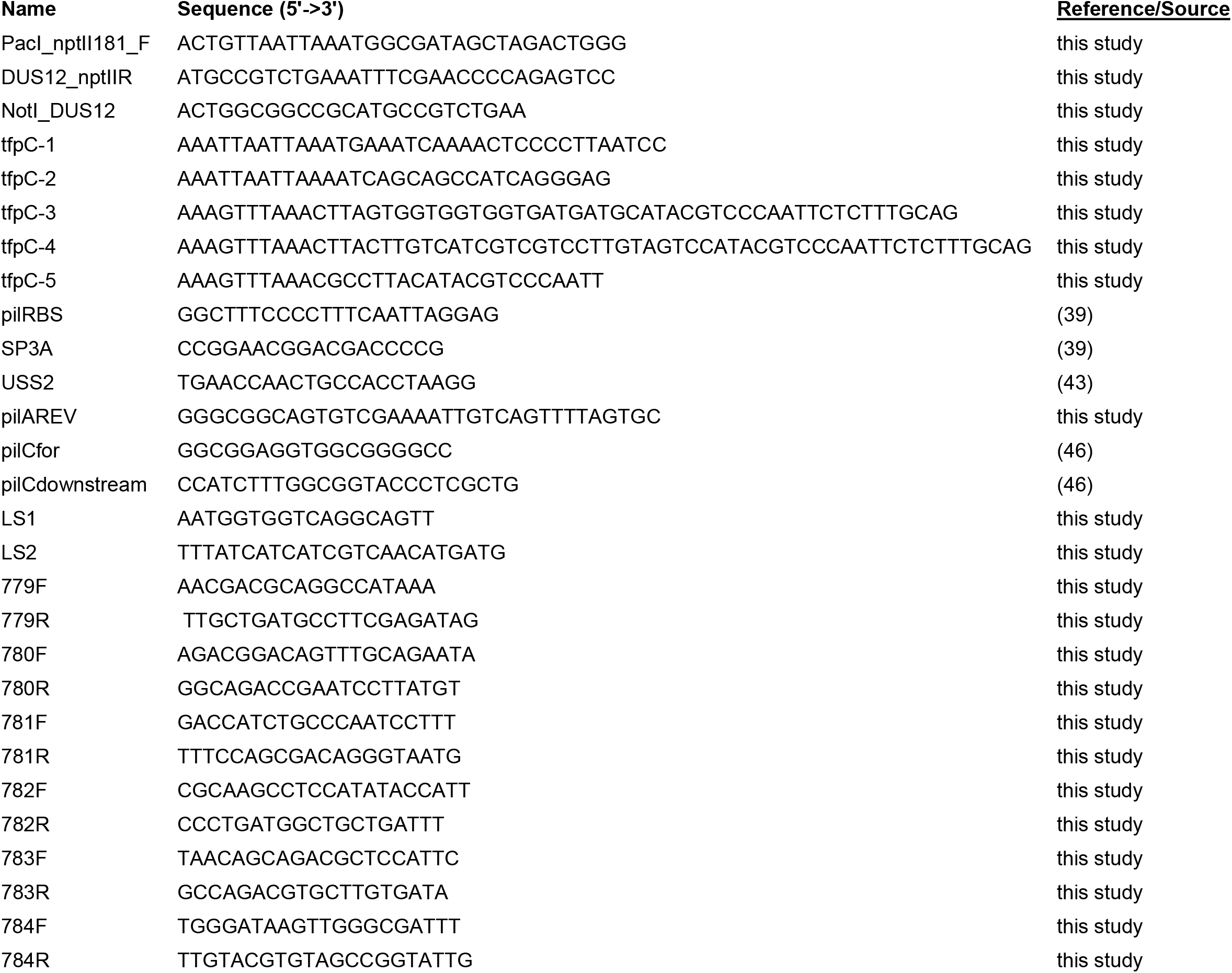
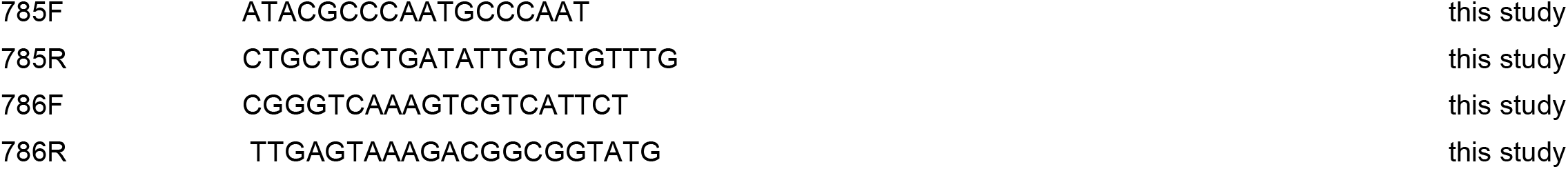
Oligonucleotides

**Table 4.**
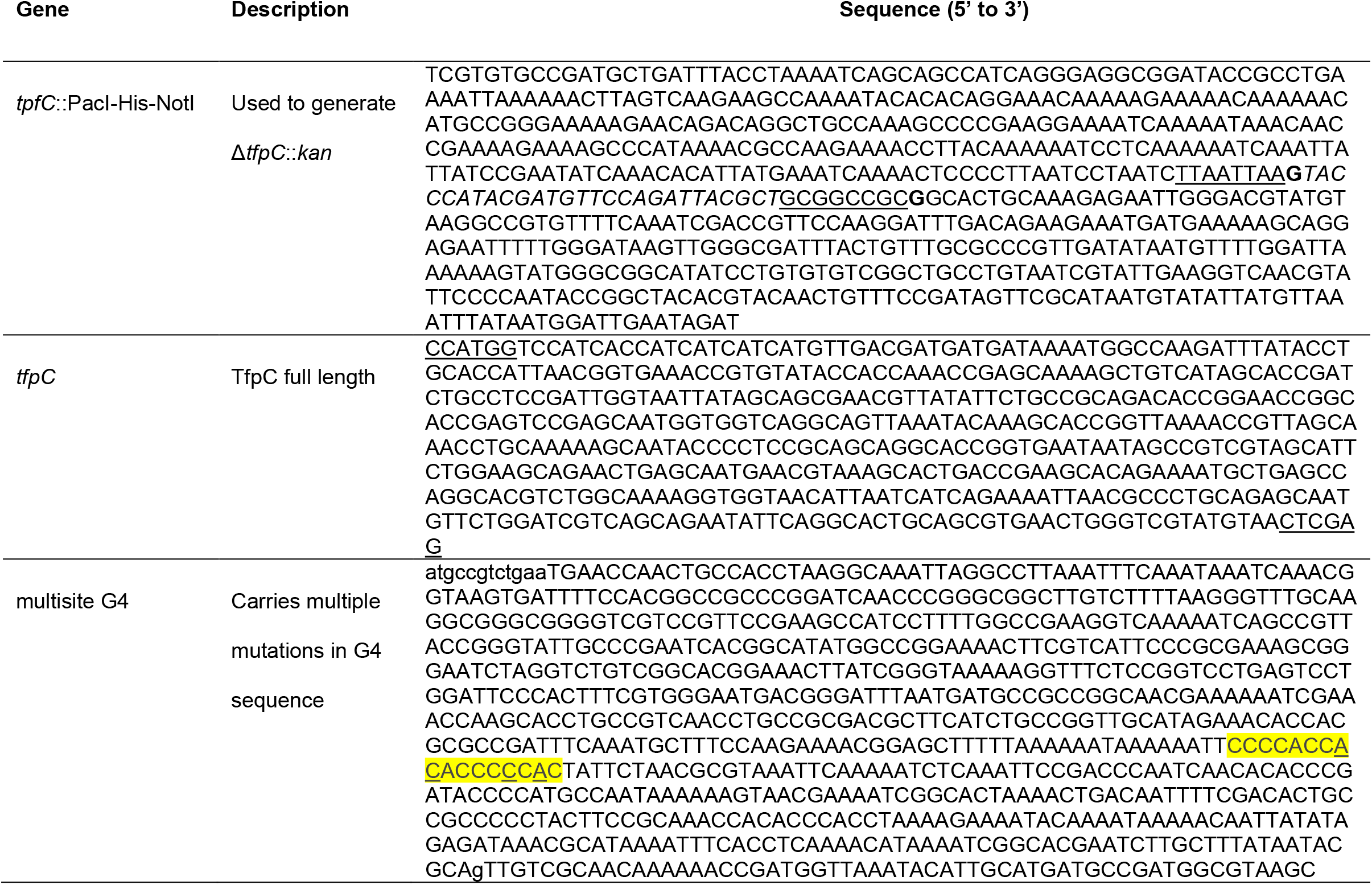
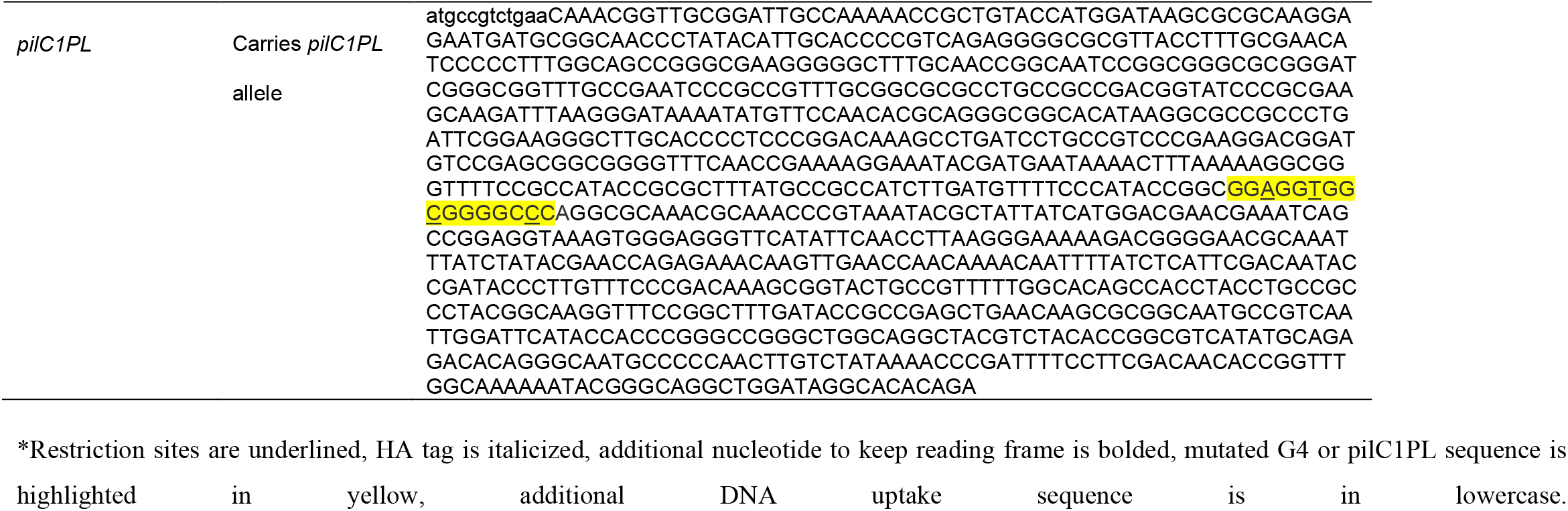
Synthetic genes.

### Constructing the parental strain N-1-60

The pilin was unable to undergo antigenic variation due to four mutations (WT: 5’-CCC CAC CCA ACC CAC CC-3’, multisite G4 mutant: 5’-CCC CAC C<u>AC</u> ACC C<u>C</u>C <u>A</u>C-3’ from Lauren Prister) in the guanine quadruplex site upstream of the *pilE* gene (40, 41). This mutant G4 sequence was introduced by synthesizing a ∼800 bp gBlock (Integrated DNA Technologies). The gBlock consisted of a DUS12 sequence and the multisite G4 substitutions flanked by regions of homology to the G4-*pilE* locus (479 bp on the 5’ end and 303 bp on the 3’ end). This DNA was used to spot transform FA1090. Several dilutions of the transformation reaction were spread onto GCB agar plates without antibiotics and grew for 41.5 hours at 37 °C in the presence of 5% CO_2_. Colonies that had a piliated colony morphology (domed surface and no blebbing from the edges) were chosen and re-streaked to confirm the piliated colony morphology. Cells that successfully recombined the multisite G4 mutations were identified by screening the piliated clones in pools by PCR. Briefly, clones were individually stored in glycerol at −80 °C and pools of 10 clones were tested by using a primer multimutG4_2 that only anneals to G4 sites that carried the desired mutations paired with RTG4-3R (42) that binds in the beginning of the *pilE* locus. Positive pools were repeated by PCR using individual clones as templates and then the promoter was amplified and sequenced with USS2 (43) and pilAREV, the *pilE* locus was amplified and sequenced with PilRBS and SP3A (39). This strain was the recipient in a transformation reaction with an approximately 950 bp gBlock (synthesized by Integrated DNA Technologies) carrying a DUS12 sequence and *pilC1*_PL_ allele (44, 45) that maintains the *pilC1* gene in a phase “on” conformation, which was flanked by 471 bp and 463 bp of homology on the 5’ and 3’ of DUS12-*pilC1*_PL_, respectively, to the *pilC1* locus. Dilutions of the transformation was grown on GCB plates and grown for 63.5 hours at 37 °C in the presence of 5% CO_2_. PCR was used to screen pools of clones that formed non-blebbing, piliated colonies before individual clones were confirmed by amplifying and sequencing *pilC1*_PL_ using pilCfor and pilCdownstream primers (46). The resultant strain is FA1090 multisite G4 mutant 1-81-S2 *pilE* variant *pilC1*_PL_ (N-1-60).

### NGO0783/Δ*tfpC mutant* construction

An approximately 650 base pair fragment containing 270 bases upstream of the *tfpC* open reading frame, the first 30 bases of TfpC, a PacI restriction site, HA tag, NotI restriction site, the last 60 bp of *tfpC* and 99 bp downstream of *tfpC*, which included a 12-mer DNA uptake sequence (DUS12) was synthesized and cloned into pTwist-Amp-MC vector by Twist Biosciences (*tfpC*::PacI-His-NotI, Table 5). A PacI- and DUS12 NotI-flanked *nptII* fragment from pBSL86 (ATCC) was generated by two PCRs: first using primers PacI_nptII181_F and DUS12_nptIIR and a second PCR to include the NotI restriction site using primers PacI_nptII181_F and NotI_DUS12. This fragment was introduced into the PacI- and NotI-digested plasmid from Twist Biosciences in between the upstream and downstream sequences of *ngo783* and in-frame. This plasmid pTwist-Δ*ngo783*::*kan* was used to spot transform several *N. gonorrohoeae* parent strains to generate Δ*tfpC* strains. Transformants were selected on GCB Kan and checked by diagnostic PCR and sequencing.

### Transformation efficiency assay

The efficiency of *N. gonorrhoeae* transformation was performed using a protocol similar to (47), except 50 ng of pSY6 DNA was used instead of 150 ng. After 20 minute incubation of the cells and DNA at 37 °C, 1 U DNase I was added to the transformation reactions and incubated for 10 minutes at 37 °C. Transformation efficiencies are reported as the mean of five independent experiments.

### Construction of tagged TfpC

The *tfpC* ORF was PCR amplified from FA1090 genomic DNA using the following primer pairs: tfpC-1 and tfpC-3 (his-tag); tfpC-1 and tfpC-4 (flag-tag); tfpC-1 and tfpC-5 (no tag). The *tfpC* fragment included the ORF and 244bp upstream of the ORF was also PCR amplified using the primer pairs as follows: tfpC-2 and tfpC-3 (His-tag); tfpC-2 and tfpC-5 (no tag). The PCR products were column purified using a PCR purification kit (Qiagen), cut by Pac1 and Pme1, and cloned into Pac1/Pme1 digested pGCC4 (48) or pGCC2 (49, 50) vector, respectively.

The resulting isopropyl-d-1-thiogalactopyranoside (IPTG)-inducible pGCC4 construct (1-2 mg) was spot-transformed into the parent (N-1-60) and Δ*tfpC* (N-3-3). The IPTG-inducible pGCC4 construct was also used to transform FA1090 1-81-S2 *recA6* Myc-tagged *pilE* (K-16-47) and the isogenic Δ*tfpC* mutant (Q115) and FA1090 1-81-S2 Myc-tagged *pilE* (Q155) and the isogenic Δ*tfpC* mutant (Q165). Strain Q155 was constructed by using a Myc-tagged *pilE* plasmid construct to transform FA1090 (32). The pGCC2 construct was spot transformed into the parent (N-1-60) and Δ*tfpC* (N-3-3). The transformants were selected on GCB with Erm and sequence confirmed.

### *pilT* mutant construction

*pilT* mutants were constructed by using 1 μg of FA1090 Δ*pilT*::*erm* genomic DNA (51) in spot transformations into the parent FA1090 1-81-S2 *myc*-tagged *pilE* (Q155) and the isogenic Δ*tfpC* mutant (Q165) and selected on GCB Erm plates.

### Western blot analysis

Colonies grown on GCB with 0.0, 0.025 mM, or 0.1 mM IPTG for 22 hours were swabbed into PBS buffer and the resuspensions were directly protein quantitated using a Pierce BCA protein assay kit (Thermo). 25 μg total protein per lane was run on a 4-15% SDS-PAGE (BioRad) at 150 V and transferred to immobilon-P membranes at 250 mA. A replicate gel was run and stained with Coomaisie Brilliant Blue to analyse total protein loading per lane. The blot was blocked in 5% nonfat milk in TBST (TBS+ 0.1% Tween 20) overnight. Anti-c-Myc antibody (Sigma) or Anti-Flag antibody (Rockland) diluted 3000x in TBST to detect the Myc-tagged PilE or Flag-tagged TfpC on a shaker for 1 hour at room temperature, respectively. The blot was washed 6 times with TBST for 5 minutes each and then incubated with 20,000x diluted secondary antibody peroxidase-conjugated goat anti-rabbit IgG (H+L) (Jackson Immuno Research) for 1 hour. After secondary antibody binding and subsequently washing, the blot was analyzed using an ECL Prime detection kit (GE Healthcare). After ECL detection of Myc-PilE, the same blot was washed 5 x 10 minutes in TBS-T using a large volume of wash buffer, blocked for 1 hour and immuno-detected using Anti-RecA (*E. coli*) antibody (1000x dilution) (gift from Mike Cox, (52)) and analyzed using an ECL Prime detection kit (GE Healthcare). Densitometry was performed using ImageJ (https://imagej.nih.gov/ij/)

### Immuno-Transmission Electron Microscopy

For analysis of piliation on strains grown on solid medium, immunoelectron microscopy was performed as described previously (53). Briefly, Formvar/carbon-coated copper grids were used to lift cells directly from 18 h old colonies and fixed for 15 min by adding a drop (17 ml) of 0.2% glutaraldehyde and 4% paraformaldehyde in Dulbecco’s PBS (DPBS; Fisher) onto the grids. The grids were washed 3 times with 1% bovine serum albumin (BSA; Sigma) in DPBS and blocked in 0.1% gelatin (Aurion, Inc.) in DPBS for 30 minutes. The grids were washed once with 1% BSA in DPBS and incubated with a 1:10 dilution of rabbit anti-c-Myc antibody (Sigma) for 1 h. Grids were washed three times with 1% BSA in DPBS and incubated with 0.1% gelatin in PBS for 30 minutes. The grids were washed once with BSA in DPBS and incubated with goat anti-rabbit IgG antibody conjugated to 12-nm gold particles (1:20 dilution; Jackson Immunolabs) for 1 h. Grids were washed five times in water for 3 minutes each. The grids were negatively stained with 1% uranyl acetate for 1 min. All washes and incubations were 17 ml and performed at room temperature. The liquid on the grids after each step was carefully wicked away using a Whatman paper. Grids were viewed using a FEI Tecnai Spirit G2 transmission electron microscope (TEM).

### Imaging of Pilus-dependent colony morphology

Representative colonies after 22 h growth on solid medium were observed and recorded using a Nikon SMZ-10A stereomicroscope and a Nikon digital sight camera.

## Quantitative RT-PCR

Overnight colonies on plain GCB plates were resuspended in GCB with 5 mM sodium bicarbonate and adjusted to OD600 ∼0.15, grown at 37 °C on a rotor for 3 hours, and treated with different concentrations of IPTG for 1 h. The cells were treated with 2 vol of RNA protect Bacteria Reagent (Qiagen) and then collected by centrifuge at 4000 rpm for 5 minutes. Total RNA was isolated using a RNeasy Mini Kit (Qiagen) and treated with RQ1 DNase (Promega) to remove genomic contamination. The quantitative RT-PCR was performed as described before (54). The 783f and 783r primer pair was used to determine the expression of TfpC with increasing IPTG concentrations. The following primer pairs were used to detect the effect of the Kan insertion into Δ*tfpC* on the surrounding genes in the operon: 1) 779f and 779r; 2) 780f and 780r; 3) 781f and 781r; 4) 782f and 782r; 5) 784f and 784r; 6) 785f and 785r; 7) 786f and 786r (Table 4).

### Cloning, expression and purification for NMR

DNA encoding full-length TfpC (residues 1–147), minus the region encoding the N-terminal periplasmic signal sequences, was synthesized by Synbio Technologies and cloned into pET28b vector using NcoI and XhoI restriction sites (Table 5). A variant encoding N-terminally truncated TfpC (TfpC-CTD; residues 52–147) was created by deletion PCR with primers LS1/2 (Table 4) using the pET28b*tfpC* plasmid as a template. Expression was carried out in *E. coli* BL21 (DE3) cells (New England Biolabs), where cells were grown in the presence of 50 μg/ml kanamycin at 37°C in M9 minimal media supplemented with ^15^NH_4_ Cl (Sigma). Expression was induced with 0.5 mM IPTG at an OD_600nm_ of 0.6 and cells were harvested after growth overnight at 18°C. Cells were resuspended in 20 mM Tris–HCl pH 8, 200 mM NaCl, lysed by sonication and purified using nickel affinity chromatography (Qiagen). Samples were then gel filtered using a Superdex 200 column (GE Healthcare) equilibrated in 20 mM Tris–HCl pH 8, 200 mM NaCl.

### NMR spectroscopy

NMR measurements were performed on 0.25 mM ^15^N-labelled samples of TfpC and TfpC-CTD in 50 mM NaPO_4_ pH 6.0, 100 mM NaCl, 1 mM tris(2-carboxyethyl)phosphine, 10% D2O or 50 mM NaPO_4_ pH 6.0, 100 mM NaCl, 10% D_2_O, respectively. 2D ^1^H-^15^N HSQC experiments were recorded with 32 scans at 298 K on a Bruker Avance III HD 700 spectrometer, equipped with TCI cryoprobe. Data were processed using NMRpipe (55) and analysed using NMRviewJ (56).

### Structural modelling

Signal peptide analysis was carried out using the SIGNALP (57) and secondary structure and domain analysis was performed using PSIPRED (58). Co-evolution analysis of mature TfpC (residues 1 to 147) was carried out using the EVcouplings Python framework (34), using default parameters. 1065 homologous sequences were identified and used in the initial alignment stage (effective sequences to protein length ratio of 5.15) and yielded 77 strong evolutionary couplings. These couplings were then used as inter-residue distance restraints to guide modelling of the TfpC structure, and within the EVcouplings Python framework. Models where the N-terminal region was folded back into the C-terminal region were discarded. The final model had a ranking score of 0.75 and was representative of the highest cluster of models.

## Acknowledgements

We thank Kyle Obergefell for providing the Myc-tagged pilin strain and Pamela Shaw for Bioinformatic support. This work was supported by Northwestern University’s NUSeq Core Facility, the Northwestern University’s Center for Advanced Microscopy (with a Cancer Center Support Grant - NCI CA060553), and the Centre for Biomolecular Spectroscopy at King’s College London for NMR access [funded by the Wellcome Trust and British Heart Foundation (ref. 202767/Z/16/Z and IG/16/2/32273 respectively)] for technical assistance. LH, SY and HSS were supported by NIH/NIAID grant R37 AI033493. LS and SR were supported by Leverhulme Trust grant RPG-2017-222 and MRC grant MR/R017662/1, respectively, awarded to JAG.

## References

1. Shaughnessy J, Ram S, Rice PA. 2019. Biology of the Gonococcus: Disease and Pathogenesis. Methods Mol Bio 1 1997:1–27.

2. Unemo M, Seifert HS, Hook EW, 3rd, Hawkes S, Ndowa F, Dillon JR. 2019. Gonorrhoea. Nat Rev Dis Primers 5:79.

3. McCarty EJ, Dinsmore WW. 2013. Important treatment change for Neisseria gonorrhoea. J Forensic Leg Med 20:181.

4. McCallum M, Burrows LL, Howell PL. 2019. The Dynamic Structures of the Type IV Pilus. Microbiol Spectr 7.

5. Denise R, Abby SS, Rocha EPC. 2020. The Evolution of Protein Secretion Systems by Co-option and Tinkering of Cellular Machineries. Trends in Microbiology 28:372–386.

6. Kellogg DS, Jr., Peacock WL, Deacon WE, Brown L, Pirkle CI. 1963. *Neisseria gonorrhoeae*. I. Virulence genetically linked to clonial variation. Journal of Bacteriology 85:1274–1279.

7. Swanson J, Robbins K, Barrera O, Corwin D, Boslego J, Ciak J, Blake M, Koomey JM. 1987. Gonococcal pilin variants in experimental gonorrhea. J Exp Med 165:1344–57.

8. Brinton CC, Jr., Wood SW, Brown A. 1982. The development of a neisserial pilus vaccine for gonorrhea and meningococcal meningitis, p 140-159. In Weinstein L, Fields BN (ed), Seminars in Infectious Disease. Thieme-Stratton, New York.

9. Pelicic V. 2008. Type IV pili: e pluribus unum? Mol Microbiol 68:827–37.

10. Merz AJ, So M. 2000. Interactions of pathogenic neisseriae with epithelial cell membranes. Annu Rev Cell Dev Biol 16:423–57.

11. Anderson MT, Byerly L, Apicella MA, Seifert HS. 2016. Seminal plasma promotes Neisseria gonorrhoeae aggregation and biofilm formation. J Bacteriol doi: 10.1128/JB.00165-16.

12. Yasukawa K, Martin P, Tinsley CR, Nassif X. 2006. Pilus-mediated adhesion of Neisseria meningitidis is negatively controlled by the pilus-retraction machinery. Mol Microbiol 59:579–89.

13. Winther-Larsen HC, Koomey M. 2002. Transcriptional, chemosensory and cell-contact-dependent regulation of type IV pilus expression. Curr Opin Microbiol 5:173–8.

14. Stohl EA, Dale EM, Criss AK, Seifert HS. 2013. Neisseria gonorrhoeae metalloprotease NGO1686 is required for full piliation, and piliation is required for resistance to H2O2-and neutrophil-mediated killing. MBio 4.

15. Freitag NE, Seifert HS, Koomey M. 1995. Characterization of the *pilF-pilD* pilus-assembly locus of *Neisseria gonorrhoeae*. Mol Microbiol 95:575–586.

16. Winther-Larsen HC, Hegge FT, Wolfgang M, Hayes SF, van Putten JP, Koomey M. 2001. Neisseria gonorrhoeae PilV, a type IV pilus-associated protein essential to human epithelial cell adherence. Proc Natl Acad Sci U S A 98:15276–81.

17. Wolfgang M, van Putten JP, Hayes SF, Koomey M. 1999. The *comP* locus of *Neisseria gonorrhoeae* encodes a type IV prepilin that is dispensable for pilus biogenesis but essential for natural transformation. Mol Microbiol 31:1345–57.

18. Martin PR, Hobbs M, Free PD, Jeske Y, Mattick JS. 1993. Characterization of pilQ, a new gene required for the biogenesis of type 4 fimbriae in Pseudomonas aeruginosa. Molecular Microbiology 9:857–868.

19. Bitter W, Koster M, Latijnhouwers M, de Cock H, Tommassen J. 1998. Formation of oligomeric rings by XcpQ and PilQ, which are involved in protein transport across the outer membrane of Pseudomonas aeruginosa. Mol Microbiol 27:209–19.

20. Jonsson AB, Nyberg G, Normark S. 1991. Phase variation of gonococcal pili by frameshift mutation in pilC, a novel gene for pilus assembly. EMBO J 10:477–88.

21. Rahman M, Kallstrom H, Normark S, Jonsson AB. 1997. PilC of pathogenic *Neisseria* is associated with the bacterial cell surface. Molecular Microbiology 25:11–25.

22. Drake SL, Sandstedt SA, Koomey M. 1997. PilP, a pilus biogenesis lipoprotein in *Neisseria gonorrhoeae*, affects expression of PilQ as a high-molecular-mass multimer. Mol Microbiol 23:657–68.

23. Rudel T, Scheurerpflug I, Meyer TF. 1995. Neisseria PilC protein identified as type-4 pilus tip-located adhesin. Nature 373:357–359.

24. Balasingham SV, Collins RF, Assalkhou R, Homberset H, Frye SA, Derrick JP, Tonjum T. 2007. Interactions between the lipoprotein PilP and the secretin PilQ in Neisseria meningitidis. J Bacteriol 189:5716–27.

25. Winther-Larsen HC, Wolfgang M, Dunham S, van Putten JP, Dorward D, Lovold C, Aas FE, Koomey M. 2005. A conserved set of pilin-like molecules controls type IV pilus dynamics and organelle-associated functions in Neisseria gonorrhoeae. ol Microbiol 56:903–17.

26. Brossay L, Paradis G, Fox R, Koomey M, Hebert J. 1994. Identification, localization, and distribution of the PilT protein in Neisseria gonorrhoeae. Infection & Immunity 62:2302–2308.

27. Park HS, Wolfgang M, Koomey M. 2002. Modification of type IV pilus-associated epithelial cell adherence and multicellular behavior by the PilU protein of Neisseria gonorrhoeae. Infect Immun 70:3891–903.

28. Stohl EA, Chan YA, Hackett KT, Kohler PL, Dillard JP, Seifert HS. 2012. Neisseria gonorrhoeae virulence factor NG1686 is a bifunctional M23B family metallopeptidase that influences resistance to hydrogen peroxide and colony morphology. J Biol Chem 287:11222–33.

29. Long CD, Hayes SF, van Putten JP, Harvey HA, Apicella MA, Seifert HS. 2001. Modulation of gonococcal piliation by regulatable transcription of *pilE*. J Bacteriol 183:1600–9.

30. Helaine S, Carbonnelle E, Prouvensier L, Beretti JL, Nassif X, Pelicic V. 2005. PilX, a pilus-associated protein essential for bacterial aggregation, is a key to pilus-facilitated attachment of Neisseria meningitidis to human cells. Mol Microbiol 55:65–77.

31. Carbonnelle E, Helaine S, Nassif X, Pelicic V. 2006. A systematic genetic analysis in Neisseria meningitidis defines the Pil proteins required for assembly, functionality, stabilization and export of type IV pili. Mol Microbiol 61:1510–22.

32. Obergfell KP. 2017. The Role of the Type IV Pilus Complex in DNA Transformation in Neisseria gonorrhoeae Northwestern University, Ann Arbor.

33. Marks DS, Colwell LJ, Sheridan R, Hopf TA, Pagnani A, Zecchina R, Sander C. 2011. Protein 3D Structure Computed from Evolutionary Sequence Variation. PLOS ONE 6:e28766.

34. Hopf TA, Green AG, Schubert B, Mersmann S, Schärfe CPI, Ingraham JB, Toth-Petroczy A, Brock K, Riesselman AJ, Palmedo P, Kang C, Sheridan R, Draizen EJ, Dallago C, Sander C, Marks DS. 2018 Python framework for coevolutionary sequence analysis. Bioinformatics 35:1582–1584.

35. Baarda BI, Zielke RA, Le Van A, Jerse AE, Sikora AE. 2019. Neisseria gonorrhoeae MlaA influences gonococcal virulence and membrane vesicle production. PLOS Pathogens 15:e1007385.

36. Reinhold-Hurek B, Hurek T. 2000. Reassessment of the taxonomic structure of the diazotrophic genus Azoarcus sensu lato and description of three new genera and new species, Azovibrio restrictus gen. nov., sp. nov., Azospira oryzae gen. nov., sp. nov. and Azonexus fungiphilus gen. nov., sp. nov. International Journal of Systematic and Evolutionary Microbiology 50:649–659.

37. Wolfgang M, van Putten JP, Hayes SF, Dorward D, Koomey M. 2000. Components and dynamics of fiber formation define a ubiquitous biogenesis pathway for bacterial pili. Embo J 19:6408–18.

38. Obergfell KP, Schaub RE, Priniski LL, Dillard JP, Seifert HS. 2018. The low-molecular-mass, penicillin-binding proteins DacB and DacC combine to modify peptidoglycan cross-linking and allow stable Type IV pilus expression in Neisseria gonorrhoeae. Molecular Microbiology 109:135–149.

39. Seifert HS, Wright CJ, Jerse AE, Cohen MS, Cannon JG. 1994. Multiple gonococcal pilin antigenic variants are produced during experimental human infections. J Clin Invest 93:2744–9.

40. Hagblom P, Segal E, Billyard E, So M. 1985. Intragenic recombination leads to pilus antigenic variation in Neisseria gonorrhoeae. Nature 315:156–8.

41. Criss AK, Kline KA, Seifert HS. 2005. The frequency and rate of pilin antigenic variation in Neisseria gonorrhoeae. Mol Microbiol 58:510–9.

42. Prister LL, Ozer EA, Cahoon LA, Seifert HS. 2019. Transcriptional initiation of a small RNA, not R-loop stability, dictates the frequency of pilin antigenic variation in Neisseria gonorrhoeae. Mol Microbiol 112:1219–1234.

43. Cahoon LA, Seifert HS. 2009. An alternative DNA structure is necessary for pilin antigenic variation in Neisseria gonorrhoeae. Science 325:764–7.

44. Cheng Y, Johnson MD, Burillo-Kirch C, Mocny JC, Anderson JE, Garrett CK, Redinbo MR, Thomas CE. 2013. Mutation of the conserved calcium-binding motif in Neisseria gonorrhoeae PilC1 impacts adhesion but not piliation. Infect Immun 81:4280–9.

45. Anderson MT, Byerly L, Apicella MA, Seifert HS. 2016. Seminal Plasma Promotes Neisseria gonorrhoeae Aggregation and Biofilm Formation. J Bacteriol 198:2228–35.

46. Rotman E, Seifert HS. 2015. Neisseria gonorrhoeae MutS affects pilin antigenic variation through mismatch correction and not by pilE guanine quartet binding. J Bacteriol 197:1828–38.

47. Obergfell KP, Seifert HS. 2016. The Pilin N-terminal Domain Maintains Neisseria gonorrhoeae Transformation Competence during Pilus Phase Variation. PLoS Genet 12:e1006069.

48. Mehr IJ. 1998. Pathways of Homologous Recombination Participate in Neisseria gonorrhoeae Pilin Antigenic Variation, DNA Transformation, and DNA RepairNorthwestern University, Evanston, IL.

49. Zhang Y, Heidrich N, Ampattu BJ, Gunderson CW, Seifert HS, Schoen C, Vogel J, Sontheimer EJ. 2013. Processing-independent CRISPR RNAs limit natural transformation in Neisseria meningitidis. Mol Cell 50:488–503.

50. Mehr IJ, Long CD, Serkin CD, Seifert HS. 2000. A homologue of the recombination-dependent growth gene, rdgC, is involved in gonococcal pilin antigenic variation. Genetics 154:523–32.

51. Long CD, Madraswala RN, Seifert HS. 1998. Comparisons between colony phase variation of *Neisseria gonorrhoeae* FA1090 and pilus, pilin, and S-pilin expression. Infection & Immunity 66:1918–27.

52. Stohl EA, Seifert HS. 2001. The *recX* gene potentiates homologous recombination in *Neisseria gonorrhoeae*. Mol Microbiol 40:1301–10.

53. Long CD, Tobiason DM, Lazio MP, Kline KA, Seifert HS. 2003. Low-level pilin expression allows for substantial DNA transformation competence in Neisseria gonorrhoeae. Infect Immun 71:6279–91.

54. Anderson MT, Seifert HS. 2013. Phase variation leads to the misidentification of a Neisseria gonorrhoeae virulence gene. PLoS One 8:e72183.

55. Delaglio F, Grzesiek S, Vuister GW, Zhu G, Pfeifer J, Bax A. 1995. NMRPipe: a multidimensional spectral processing system based on UNIX pipes. J Biomol NMR 6:277–93.

56. Johnson BA, Blevins RA. 1994. NMR View: A computer program for the visualization and analysis of NMR data. Journal of Biomolecular NMR 4:603–614.

57. Almagro Armenteros JJ, Tsirigos KD, Sønderby CK, Petersen TN, Winther O, Brunak S, von Heijne G, Nielsen H. 2019. SignalP 5.0 improves signal peptide predictions using deep neural networks. Nature Biotechnology 37:420–423.

58. Buchan DWA, Jones DT. 2019. The PSIPRED Protein Analysis Workbench: 20 years on. Nucleic Acids Research 47:W402–W407.

59. Mehr IJ, Seifert HS. 1998. Differential roles of homologous recombination pathways in *Neisseria gonorrhoeae* pilin antigenic variation, DNA transformation, and DNA repair. Molecular Microbiology 30:697–710.

60. Haas R, Schwarz H, Meyer TF. 1987. Release of soluble pilin antigen coupled with gene conversion in *Neisseria gonorrhoeae*. Proceedings of the National Academy of Sciences of the United States of America 84:9079–9083.

